# Screening antifungal and antioxidant activity of macroalgae from SE Spain highlights the invader *Rugulopteryx okamurae*

**DOI:** 10.64898/2026.04.07.716908

**Authors:** M. Valverde-Urrea, C.K. Otero, M. Terradas-Fernández, F. Lopez-Moya

## Abstract

The Mediterranean Sea harbors a rich diversity of macroalgae with pharmacological potential. In this study, metabolite composition, antioxidant and antifungal activities of methanol and ethyl acetate extracts from *Rugulopteryx okamurae, Dictyota fasciola, Batophora* sp., *Codium fragile*, and *Palisada tenerrima* from the southeastern coast of Spain were evaluated. *R. okamurae, Batophora* sp. and *C. fragile* are non-native. All extracts exhibited antioxidant activity, particularly those obtained with methanol. *R. okamurae* and *Batophora* sp. showed the highest activity, inhibiting the DPPH·radical by more than 40% at 1 mg/ml. All extracts contained phenolics and flavonoids, which may contribute to the observed antioxidant activity. Moreover, the methanolic extracts of *R. okamurae* and *P. tenerrima* exhibited *in vitro* fungistatic activity against the wilt pathogen *Fusarium oxysporum* f. sp. *cubense* tropical race 4. *R. okamurae* extracts showed the strongest antifungal activity against *F. oxysporum* f. sp. *cubense* TR4, with inhibition values of 23.3% and 30.5% at doses of 10 and 20 mg/well, respectively. The methanolic *P. tenerrima* also showed notable activity (19.8% and 20.7% inhibition), whereas other extracts displayed lower effect. LC–MS/MS analysis of *R. okamurae* extract revealed a diverse metabolite profile including oxylipin-type metabolites, terpenoid-like compounds and carotenoids. Our findings highlight coastal macroalgae from SE Spain as sources of bioactive compounds and support the valorization of biomass from invaders such as *R. okamurae*.

## 1. Introduction

Marine macroalgae are a rich source of bioactive compounds with promising pharmacological potential. In recent years, growing interest has emerged in exploring algae as natural sources of antioxidants and antifungals, for their unique secondary metabolites profiles and sustainable accessibility (Batista et al., 2009; Mayer et al., 2009). Among these, species inhabiting the southeastern coast of Spain—including both native and invasive taxa—have drawn attention for their abundant biomass and potential for biotechnological applications.

Invasive macroalgae such as *Rugulopteryx okamurae, Codium fragile* or *Batophora* sp. have proliferated along the Mediterranean coast, disrupting ecosystems and local biodiversity (Sempere-Valverde et al., 2021; Terradas-Fernández et al., 2020; Terradas-Fernández et al., 2023). Beyond their ecological impacts, these alien species represent an untapped biomass resource with potential value for bioactive compound extraction. Using this biomass for obtaining antioxidant and antifungal extracts could not only generate value-added products but also contribute to mitigating their environmental impact (Alidoost et al., 2021; El Mandany et al., 2023).

Several studies have highlighted the antioxidant capacity of marine algal extracts, attributed mainly to their content of phenolic compounds, flavonoids, terpenoids, carotenoids, and other metabolites (Vidal et al., 2009; Salehi et al., 2019). Antioxidants from marine algae may play an important role in neutralizing reactive oxygen species (ROS), thereby preventing oxidative stress-induced cellular damage—a process implicated in the development of various chronic diseases including cancer and neurodegenerative disorders (Sies et al., 2017; Hajam et al., 2022).

In addition to their antioxidant potential, algal extracts have demonstrated significant antifungal activity against phytopathogens such as *Fusarium oxysporum*, including the highly aggressive f.sp.cubense tropical race 4 (TR4) responsible for Fusarium wilt in banana crops (El-Sheekh et al., 2020). Current control measures are limited, and no effective treatment exists for this pathogen, highlighting the need for alternative, sustainable antifungal agents (Tripathi et al., 2009; Martínez-Solórzano et al., 2020). Algal secondary metabolites may offer promising antifungal activities, as demonstrated in previous in vitro studies (Mostafa et al., 2022; Elshikh and Al Farraj, 2024).

Extraction methods and solvent polarity are known to influence the recovery of bioactive compounds from algae, with polar solvents such as methanol showing greater efficacy in extracting phenolic and flavonoid compounds, thereby enhancing antioxidant and antifungal activities (Monteiro et al., 2020; Mendes et al., 2013).

Given the increasing demand for natural antioxidants and antifungals, coupled with the environmental need to manage invasive algal species, the valorisation of marine macroalgae from the southeastern coast of Spain represents a viable strategy for sustainable biotechnological development.

Therefore, the objective of this study is to evaluate the antioxidant and antifungal activities of methanol and ethyl acetate extracts from selected macroalgae collected from accessible coastal areas of southeastern Spain. We also aim to assess the potential of invasive species biomass for biotechnological applications, thereby contributing to both environmental management and the identification of promising algal resources for future research.

## 2. Materials and Methods

### 2.1. Collection and identification of algal species

Algal samples were cleaned of sediments and epiphytes and identified morphologically using specialized keys (Cormaci et al., 2012, 2014, 2020; Terradas-Fernández et al., 2022, 2023; Vitales et al., 2019). Molecular identification was performed following a CTAB-based DNA extraction protocol (Coat et al., 1998) with minor modifications. Briefly, freeze-ground algal tissue (liquid nitrogen) was incubated in CTAB buffer (0.1 M Tris–HCl, 0.05 M EDTA, 1.5 M NaCl, 0.05 DTT, 2% PVP, 3% CTAB) at 60 °C, followed by chloroform:isoamyl alcohol (24:1, v/v) extraction and ethanol precipitation. DNA pellets were washed with 70% ethanol, air-dried, and resuspended in nuclease-free water. DNA quality and concentration were assessed using a NanoDrop One C spectrophotometer (Thermo Fisher Scientific).

PCR amplifications were performed using VWR Red Taq DNA polymerase Master Mix in a LifeExpress thermal cycler (BIOER). For brown algae, the psbA gene was amplified using primers psbAF1 (5⍰-ATGACTGCTACTTTAGAAAGAC-3⍰) and psbAR2 (5⍰-TCATGCATWACTTCCATACCTA-3⍰) (Saunders and Moore, 2013) under a two-step cycling protocol: an initial denaturation (94 °C, 2 min), followed by 5 cycles (94 °C for 30 s, 45 °C for 30 s, 72 °C for 1 min), and 35 cycles (94 °C for 39 s, 46.2 °C for 39 s, 72 °C for 1 min), with a final extension at 72 °C for 7 min. Green algae were identified by amplification of the rbcL gene using primers rbcL B (5⍰-ATGTCACCACAAACAGAAACTAAAGCA-3⍰) and rbcL Q (5⍰-GATCTCCTTCCATACTTCACAAGC-3⍰) (Zechman, 2003), with cycling conditions of 94 °C for 3 min; 30 cycles of 94 °C for 1 min, 50 °C for 30 s, and 72 °C for 1.5 min; and a final extension at 72 °C for 7 min. For red and brown algae, the COI gene was amplified using primers GazF2 (5⍰-CCAACCAYAAAGATATWGGTAC-3⍰) and GazR2 (5⍰-GGATGACCAAARAACCAAAA-3⍰) (Lane et al., 2007), with an initial denaturation at 94 °C for 4 min, followed by 40 cycles (94 °C for 1 min, 50 °C for 30 s, 72 °C for 1 min) and a final extension at 72 °C for 7 min.

PCR products were verified on 1% agarose gels, purified using the GeneJET Gel Extraction Kit (Thermo Scientific), and sequenced by Sanger sequencing (STAB Vida, Portugal). Resulting sequences were compared against the National Center for Biotechnology Information database using BLASTn.

### 2.2. Extract preparation

Fresh algal samples were freeze-dried for 48 hours in a CHRIST Alpha 1-2 LDplus lyophilizer (Martin Christ Gefriertrocknungsanlagen GmbH, Osterode am Harz, Germany). Moisture content was calculated by weight difference. Dried samples were ground in liquid nitrogen.

Extracts were prepared by mixing 20 g of powdered algae with methanol (MeOH) or ethyl acetate (EtoAc) HPLC grade at 1:10 (w/v) ratio (Thangara et al. 2019). Extraction involved heating at 40 °C under stirring for 2 hours, followed by 48 hours stirring at 4 °C. Extracts were filtered through Whatman No.1 filter paper and 0.22 µm syringe filters (Merck KGaA, Darmstadt, Germany), evaporated at 45 °C using a Büchi R-100 rotary evaporator (Büchi Labortechnik AG, Flawil, Switzerland), and stored at -20 °C. Dried extracts were dissolved in dimethyl sulfoxide (DMSO) to a final stock concentration of 100 mg/ml. Extraction yield was calculated as:

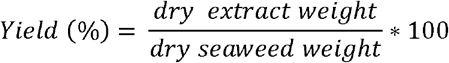

Each extraction was performed in triplicate and pooled to reduce variability.

### 2.3. Extract characterization

#### 2.3.1. Total phenolic content

Phenolic content was quantified using the Folin-Ciocalteu method (Singleton et al., 1999). Absorbance was measured at 760 nm using a Thermo Spectronic HEλIOS ε spectrophotometer (Thermo Fisher Scientific, Massachusetts, USA), expressed as µg gallic acid equivalents per mg dry extract.

#### 2.3.2. Total flavonoid content

Flavonoid content was measured by mixing 0.5 ml extract with 1 ml 10% AlCl_3_ and 0.5 ml 120 mM potassium acetate, incubating 30 min at 30 °C, and reading absorbance at 415 nm (Md. Monir et al., 2015). Results were expressed as µg quercetin equivalents per mg dry extract.

### 2.4. Antioxidant activity

Antioxidant activity was assessed using the 2,2-diphenyl-1-picrylhydrazyl (DPPH) radical scavenging assay (Kedare and Singh, 2011). A 0.1 ml volume extract (0.25, 0.50, 0.75, 1 mg/ml) was mixed with 2.9 ml 3.9 mg/100 ml DPPH in 90% methanol, incubated in the dark for 30 min at 4 °C, and absorbance measured at 517 nm. Scavenging capacity was calculated as:

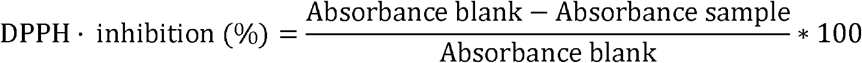

Ascorbic acid (250–1000 µg/ml) was used as positive control.

All colorimetric assays (TPC, TFC, protein content, DPPH) were performed using a Thermo Spectronic HELIOS ε spectrophotometer (Thermo Fisher Scientific, MA, USA). All experiments were carried out in triplicate.

### 2.5. Antifungal activity

Agar diffusion assays were used to evaluate antifungal activity against *Fusarium oxysporum* f. sp. *cubense* TR4 (Foc TR4). PDA plates (1% agar) were prepared with six wells containing algal extracts, DMSO controls, nystatin (17.5 µg; positive control), and an empty well (negative control). Extracts were prepared in DMSO (100 mg mL^−1^) and 100 or 200 µL were added to each well, corresponding to doses of 10 and 20 mg per well, respectively. An 8-mm mycelial plug of Foc TR4 was placed at the center of each plate, and radial growth was recorded daily for 10 days.

### 2.6. HPLC-MS/MS analysis

The compounds present in the methanol and ethyl acetate extracts were analysed by high-performance liquid chromatography coupled to tandem mass spectrometry (HPLC–MS/MS) using an Agilent 1290 Infinity system coupled to an Agilent 6550 iFunnel Q-TOF mass spectrometer. Mass axis calibration was performed by continuous infusion of a reference solution consisting of 95% acetonitrile and 5% water. Chromatographic separation was carried out using a Zorbax Eclipse Plus C18 column (2.1 × 100 mm, 1.8 μm particle size). The injection volume was 1 μL and the analysis was conducted at 4 °C over a 20 min run.

Data processing was performed using the Agilent MassHunter software suite. MassHunter Data Acquisition 10.1 was used to operate the LC/QTOF 6550 system, while MassHunter Qualitative Analysis B.10.0 together with MassHunter PCDL Manager B.08 were used for qualitative analysis of the data. Compound identification was performed by comparison of MS/MS spectra with the METLIN Agilent compound database.

### 2.7. Statistical analysis

Extraction yield was assessed with two-way ANOVA (species × solvent), while antioxidant capacity was evaluated by three-way ANOVA (species × solvent × extract concentration). Phenolic and flavonoid contents were analyzed by two-way ANOVA (species × solvent).

The antifungal activity was analyzed with generalized linear models (GLM, Gaussian family), including day and treatment as predictors and repeated measures as a random effect. Growth data at day 7 were further analyzed by one-way ANOVA (treatment). Post hoc tests included TukeyHSD for ANOVA and pairwise *t*-tests with Sidak correction (emmeans; Lenth, 2024). Model assumptions were tested with Shapiro–Wilk and Levene’s tests; GLM validation was performed with DHARMa (Hartig, 2022). If parametric assumption were not met, data were square-root transformed.

To integrate biological and chemical performance and prioritize the most promising algal extracts, a multiobjective Pareto analysis was applied. This approach identifies extracts that achieve the best overall balance between antifungal activity, antioxidant potential, and extraction yield, allowing a multidimensional comparison without imposing arbitrary weights.

The analysis considered six maximization criteria: antifungal inhibition at 10 and 20 mg/well, DPPH scavenging activity at 1 mg/ml, total phenolic and flavonoid contents (µg GAE/mg and µg QE mg, respectively), and extraction yield (%).

Each extract (species × solvent) was represented as a point in this multidimensional space and classified as Pareto-optimal when no other extract outperformed it simultaneously in all variables. Pareto-optimal extracts thus define the non-dominated frontier, representing the most balanced and efficient candidates for downstream fractionation and valorisation.

All analyses and graphics were performed in R v4.3.3 (R Core Team, 2023).

## 3 Results

### 3.1. Identification of algal species

*Rugulopteryx okamurae* (E.Y. Dawson) I.K. Hwang, W.J. Lee and H.S. Kim, 2009 shows a yellow– brown thallus with dichotomous branching (isotomous and anisotomous), reaching up to seven branching orders. In the apical and medial zones, the medulla appears single-layered medially and multistratified at the margins, with two to three marginal layers (Figure S1).

*Dictyota fasciola* (Roth) J.V. Lamouroux, 1809 exhibits a yellow–brown thallus, with lighter yellow coloration in apical regions. Branching occurs forming dichotomies with two to four branching orders. Axes show a spiral arrangement and acute apical terminations. Both the medulla and marginal regions are unstratified in apical and medial zones (Figure S2).

Codium fragile (Suringar) Hariot, 1889 develops a dark green siphonous plectenchymatous thallus with dichotomous branching. Utricles are green, globose, and mucronate, lacking apical teeth (Figure S3).

*Batophora* sp. J. Agardh, 1854 displays a yellow–green thallus with darker green branches. The central cylindrical axis bears verticils of 7–11 branches, showing dichotomous and trichotomous branching up to seven orders (Figure S4).

*Palisada tenerrima* (Cremades) D. Serio, M. Cormaci, G. Furnari and F. Boisset, 2010 presents a green–brown thallus that turns violet in older basal branches. Attachment occurs through a basal disc (≈2 cm diameter). Branching follows three to four orders. Cells are irregularly polyhedral and slightly parallel in arrangement, with no *corps en cerise* or secondary pit connections. Transverse sections reveal elongated palisade-like cortical cells (Figure S5).

Molecular analysis confirmed the morphological identification for all species except *C. fragile*, whose DNA could not be amplified (Table S1).

### 3.2 Chemical characterization and extraction yield of the studied algae

All species showed a water content above 70%, with significant differences among them (ANOVA, F_4_,_28_ = 12.97, p < 0.001) (Figure S6A, Table S2). *C. fragile* registered the lowest value (79.7 ± 4.98%), whereas *D. fasciola* reached the highest (94.5 ± 2.49%).

Extraction yield depended on both solvent and species (two-way ANOVA, F_4_,_20_ = 26.97, p < 0.001) (Figure S6B, Table S3). Methanolic extracts generally produced higher yields than ethyl acetate extracts. Exceptions included *Batophora* sp., where solvent had no significant effect. Among methanolic extracts, *D. fasciola* and *R. okamurae* achieved the highest yields (12.5±0.41 and 12.4±0.34), while ethyl acetate extraction favoured *R. okamurae* (10.2 ± 0.52%). *Batophora* sp. and *P. tenerrima* produced the lowest yields overall.

Total phenolic content varied significantly across species and solvents (two-way ANOVA, F_4_,_50_ = 17.04, p < 0.001) (Figure 1A, Table S4). Methanol consistently extracted higher levels of phenolics, particularly in *R. okamurae* (62.3 ± 13.7 µg GAE/mg). In contrast, *P. tenerrima* contained the lowest phenolic levels (13.7 ± 1.99 µg GAE/mg). Within ethyl acetate extracts, *D. fasciola* showed the highest phenolic concentration (47.6 ± 13.3 µg GAE/mg).

**Figure 1.**
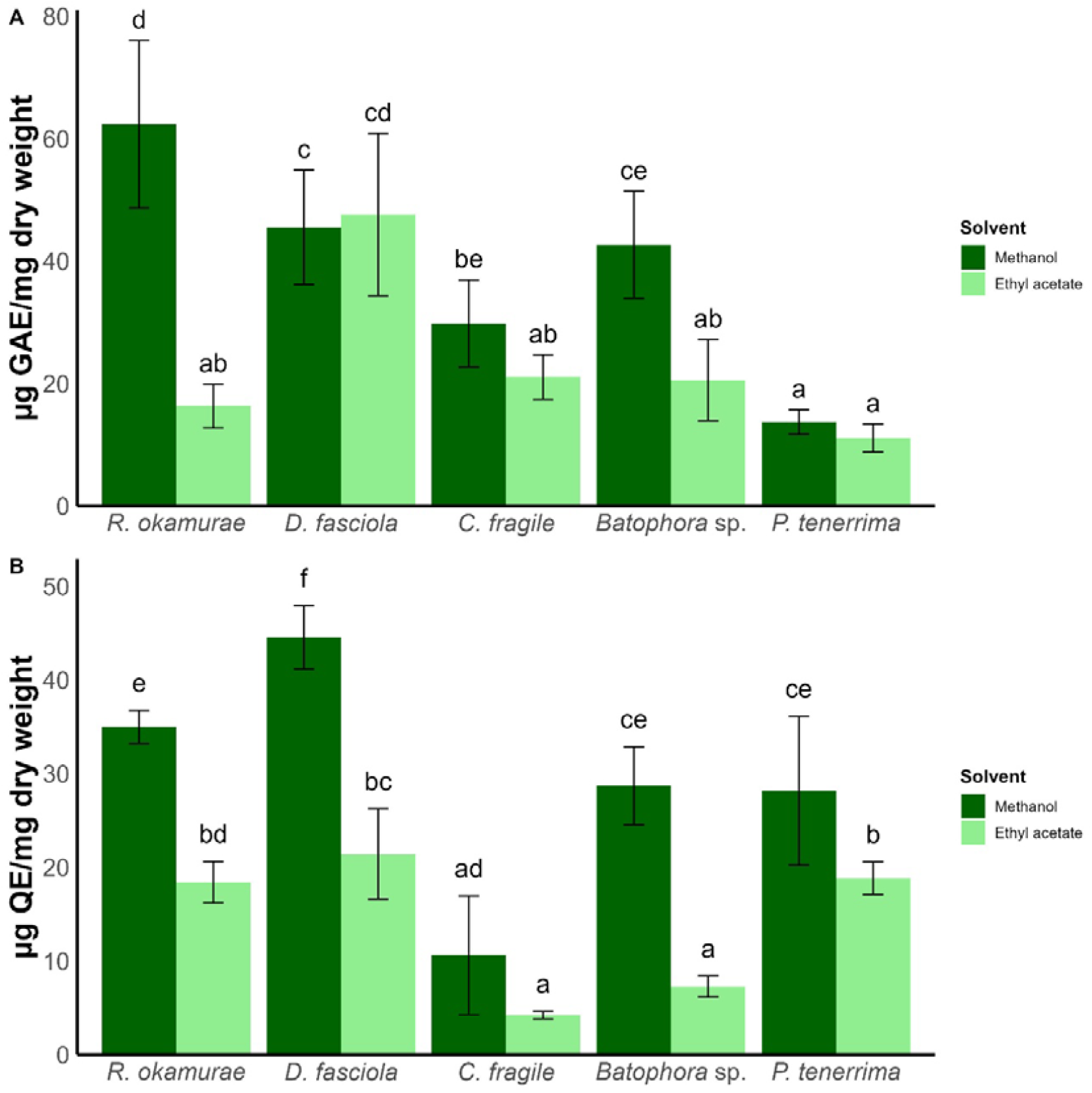
Phenolic (A) and flavonoid (B) content according to algae species and solvent used. Different letters indicate statistically significant differences (*p* < 0.05).

Flavonoid content also depended on solvent and species (two-way ANOVA, F_4_,_50_ = 9.68, p < 0.001) (Figure 1B, Table S5). Methanol extracts accumulated higher flavonoid concentrations than ethyl acetate, except in *C. fragile*, where flavonoid levels were similar in both solvents. The methanolic extract of *D. fasciola* reached the maximum (44.6 ± 3.37 µg QE/mg). Ethyl acetate extracts of *D. fasciola* and *R. okamurae* still surpassed 25 µg QE/mg, while *C. fragile* showed the lowest flavonoid content (4.24 ± 0.40 µg QE/mg).

### 3.3 Antioxidant potential of algal extracts

The antioxidant activity of the extracts was influenced by species, solvent, and extract concentration (three-way ANOVA, F_15_,_96_ = 3.07, p < 0.001) (Table S6).

For methanolic extracts (Figure 2A), antioxidant capacity is positively correlated with extract concentration, though this trend was only statistically significant for *C. fragile* (21 ± 3.38% DPPH inhibition at 0.25 mg/ml vs. 35.8 ± 2.08% at 1 mg/ml; Figure 2A). Among methanolic extracts, *Batophora* sp. exhibited the highest antioxidant activity (44.9 ± 4.89% at 1 mg/ml), closely followed by *R. okamurae* (44.68 ± 0.86% DPPH inhibition at 1 mg/ml). The remaining species showed DPPH inhibition below 40% at all tested concentrations.

**Figure 2.**
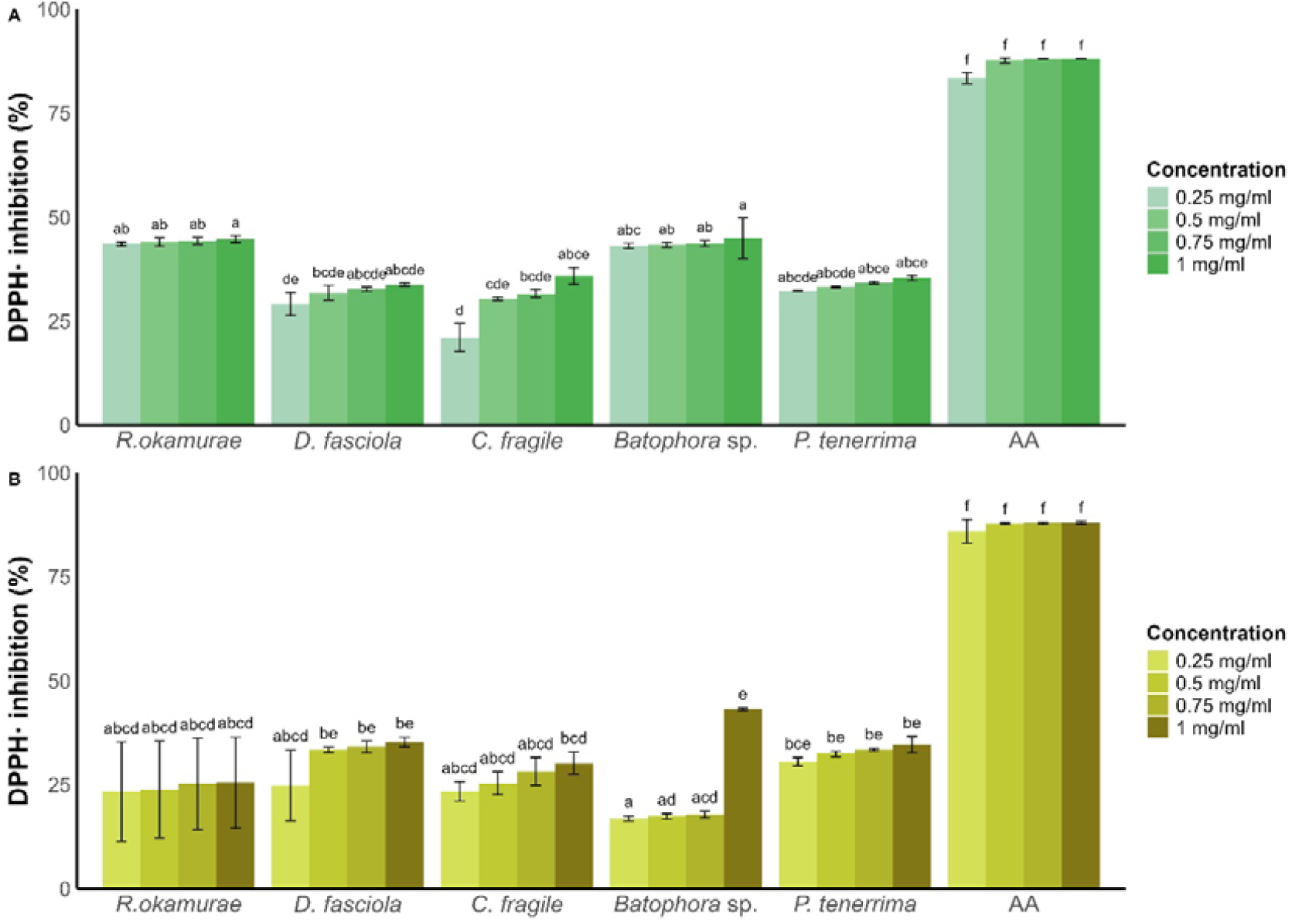
DPPH· radical scavenging activity (%) according to algae species and concentration for methanol (A) and (B) ethyl acetate extracts. AA: ascorbic acid. Different letters indicate statistically significant differences (*p* < 0.05).

For ethyl acetate extracts (Figure 2B), the highest antioxidant activity was observed for *Batophora* sp. extracts (43.19 ± 0.30% DPPH inhibition at 1 mg/ml). This extract showed a 25.32% increase in antioxidant capacity between 0.75 mg/ml and 1 mg/ml, without a clear linear dose-response pattern.

All other ethyl acetate extracts exhibited DPPH inhibition below 35%, with *R. okamurae* showing the lowest antioxidant activity (25.54 ± 10.94% at 1 mg/ml).

Comparing the antioxidant capacity at the highest concentration (1 mg/ml), methanolic extracts generally outperformed ethyl acetate extracts across all species—except for *Batophora* sp., where the ethyl acetate extract had superior antioxidant activity.

### 3.4. Antifungal Activity

Among all tested algal extracts, the methanolic extract of *R. okamurae* exhibited the most potent antifungal effect against *Fusarium oxysporum* f. sp. *cubense* TR4 (30.5±9.13 % inhibition relative to the control). A significant interaction was found between treatment and time (GLM, F_45_,_118_ = 1.58, *p* = 0.02) (Figure 3 and S7, Table S7), indicating that fungal growth was reduced over time depending on the treatment applied. From day 6 onwards, 20 mg/well of extract significantly reduced colony expansion compared to the control and even to the commercial antifungal nystatin. At day 7 (D7) (Table S8), a significant treatment effect was also detected (ANOVA, F_5_,_12_ = 6.22, *p* < 0.001), with 20 mg of extract per well reducing growth to 2.53 ± 0.20 mm, compared to 3.66 ± 0.23 mm in the control.

**Figure 3:**
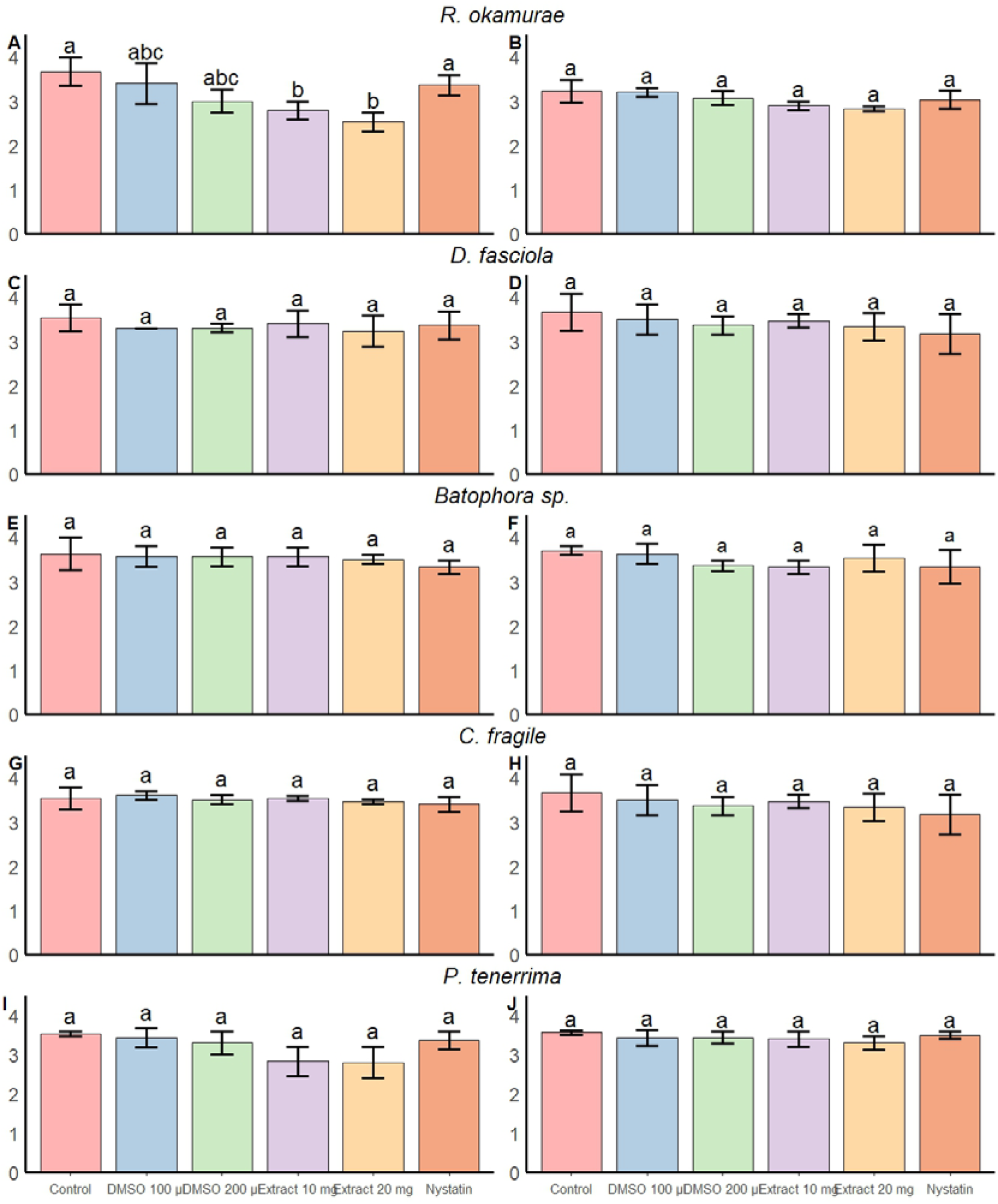
Effect of algal extracts obtained with methanol (left column) and ethyl acetate (right column) on the growth front of *Foc TR4*.Panels correspond to extracts from *R. okamurae* (A–B), *D. fasciola* (C–D), *C. fragile* (E–F), *Batophora* sp. (G–H), and *P. tenerrima* (I–J). Bars represent mean ± SD. Different letters above the bars indicate statistically significant differences between treatments (p < 0.05).

Similarly, the methanolic extract of *P. tenerrima* significantly inhibited fungal growth (GLM, F_5_,_118_ = 24.68, *p* < 0.001), with both 10 mg and 20 mg/well doses per well showing clear inhibitory effects compared to control (19.8 ± 10.4 and 20.7 ± 11.5% inhibition relative to control). However, post hoc comparisons at D7 failed to detect significant differences among treatments (ANOVA, F_5_,_12_ = 3.48, *p* = 0.03), though a reduction trend was observed (control: 3.53 ± 0.05 mm; 20 mg/well extract: 2.80 ± 0.40 mm; nystatin: 3.37 ± 0.23 mm).

Other extracts showed more limited or inconsistent effects. The ethyl acetate extract of *P. tenerrima* showed no significant inhibition despite a statistically significant GLM result (F_5_,_118_ = 3.74, *p* = 0.03). Similarly, *D. fasciola* and *Codium fragile* extracts had variable outcomes: the ethyl acetate extract of *D. fasciola* significantly reduced growth (GLM, F_5_,_117_ = 8.27, *p* < 0.0001), but this effect was not significant at D7. The methanolic extract of *C. fragile* showed a significant treatment effect (GLM, F_5_,_118_ = 5.32, *p* < 0.001), but no individual treatment reached significance in post hoc tests. In contrast, the ethyl acetate extract of *C. fragile* at 10 mg/well did reduce fungal growth significantly.

*Batophora* sp. extracts displayed a mild and inconsistent antifungal effect. The methanolic extract showed no significant inhibition at any concentration, whereas the ethyl acetate extract at 20 mg/well slightly reduced fungal growth (GLM, F_5_,_118_ = 16.86, *p* < 0.001), though the effect was less pronounced than that of nystatin.

### 3.5 Multi-criteria optimization of algal extracts through Pareto analysis

The five-dimensional Pareto analysis (criteria: antifungal inhibition at 10 and 20 mg/well, DPPH scavenging at 1 mg/ml, total phenolics, total flavonoids, and extraction yield) revealed a small subset of globally optimal extracts (Figure 4 and Table S9). In this approach, extracts are considered Pareto-optimal when no other extract performs better simultaneously across all evaluated criteria. Methanolic fractions of *R. okamurae, D. fasciola*, and *Batophora* sp. met this condition.

**Figure 4:**
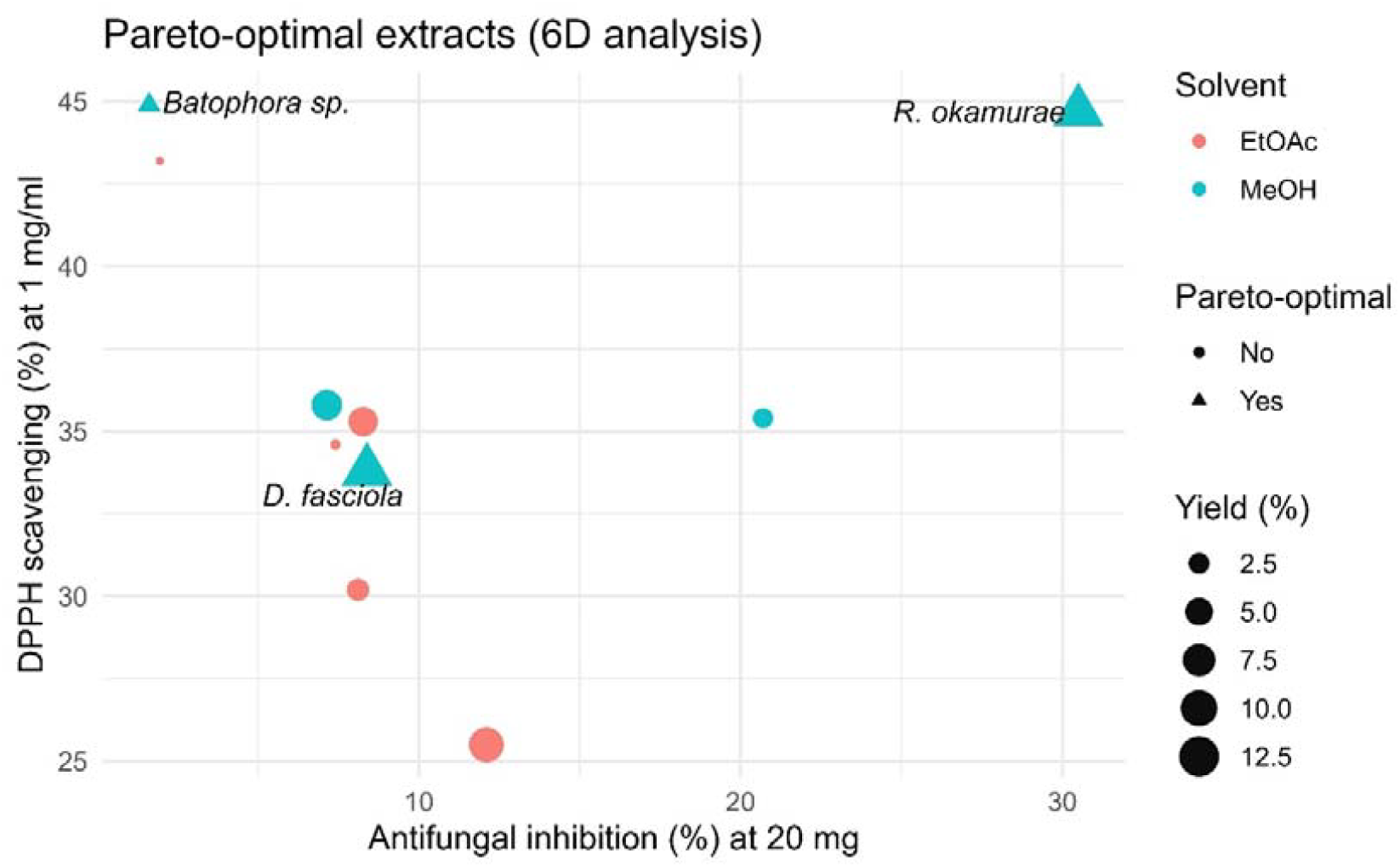
Pareto-optimal extracts (6D analysis). Scatter plot of antifungal inhibition at 20 mg/ml (X) versus DPPH scavenging at 1 mg/ml (Y). Point size denotes extraction yield (%); color indicates solvent (MeOH, EtOAc). Triangles mark Pareto-optimal extracts computed over six maximization criteria (inhibition at 10 and 20 mg/well, DPPH, total phenolics, total flavonoids, and yield). Labels show *R. okamurae, D. fasciola*, and *Batophora* sp. as globally optimal candidates.

Among them, the methanolic extract of *R. okamurae* provided the best overall compromise, combining the highest antifungal inhibition (∼30% at 20 mg/well), strong antioxidant capacity (∼45% DPPH), and the greatest extraction yield (∼12%). *D. fasciola* (methanol extract) remained optimal through a balanced profile (moderate inhibition with comparatively high phenolic/flavonoid content and mid-range DPPH), whereas *Batophora* sp. (methanol extract) was mainly driven by exceptional antioxidant metrics (∼45% DPPH) despite modest antifungal effects. All ethyl acetate extracts were dominated in the multidimensional space, underscoring the superior multifunctional performance of polar (methanolic) fractions.

### 3.6. HPLC-MS/MS extract analysis

Following the Pareto optimisation, the methanolic extract of *R. okamurae* emerged as most promising, combining high antioxidant activity, elevated phenolic content and strong antifungal effects against *Fusarium oxysporum* f. sp. *cubense* TR4. The ethyl acetate extract also showed relevant bioactivity in the Pareto analysis and was therefore included for comparative chemical profiling. To explore the chemical composition of both extracts, we used LC-MS.

Methanolic extracts showed by LC-MS a complex mixture of metabolites distributed across the retention time range (Table 10S and 11S). Accurate mass measurements and MS/MS fragmentation patterns indicated the presence of phenolic derivatives, oxylipin-type metabolites derived from C18 fatty acids, hydroxy fatty acids and terpenoid-like compounds. Several late-eluting signals matched sulfolipid derivatives commonly found in photosynthetic organisms. The analysis also revealed carotenoid pigments characteristic of brown macroalgae, including fucoxanthin and related xanthophyll derivatives.

We also found oxylipin-related compounds such as 12-oxophytodienoic acid (12-OPDA) likely from oxidation of polyunsaturated fatty acids. The chromatographic profile also showed signals consistent with fucoxanthin, a carotenoid widely reported in brown algae. In addition, a peak detected at intermediate retention time corresponded to a compound with molecular formula C_15_H_22_O_2_, consistent with a bisabolene-type sesquiterpenoid related to curcumenone-like structures.

The LC–MS profile of the ethyl acetate extract showed a broadly similar but slightly less complex metabolite pattern. The signals mainly corresponded to terpenoid-like compounds, aromatic derivatives and lipid-related metabolites. Oxylipin-type compounds derived from polyunsaturated fatty acids also appeared in this extract, including signals compatible with 12-OPDA and related oxidation products.

Several peaks detected at intermediate retention times corresponded to compounds with molecular formulas compatible with sesquiterpenoid structures, including signals consistent with bisabolane-type compounds. Carotenoid pigments characteristic of brown macroalgae, including fucoxanthin, were also detected in this extract.

Overall, the ethyl acetate extract showed a lower diversity of polar metabolites compared with the methanolic extract, which may partly explain the lower antioxidant and antifungal activities observed in the bioassays.

## 4. Discussion

This study evaluated extracts from native and invasive macroalgae from the southeastern Spanish coast. They displayed both antioxidant and antifungal activities.

Methanolic extractions consistently yielded higher recoveries than ethyl acetate, in line with previous reports showing that polar solvents, as methanol, extract efficiently algal metabolites (Nair et al., 2007; López et al., 2011). Our yields (≈2–12%) fit within the ranges reported for similar methods (Rabecca et al., 2022; Vasconcelos et al., 2017). Differences with other studies may reflect solvent polarity, time of extraction, or diverse species composition (Cho et al., 2011; Farvin and Jacobsen, 2013). The maceration method applied here also explains the lower yields compared to Soxhlet or other optimized techniques (Tambun et al., 2021).

Phenolic and flavonoid contents varied among species. In *R. okamurae* and *D. fasciola*, values ranged from 17–60 µg GAE/mg, below the higher levels reported for other brown algae when using optimized methods (Getachew et al., 2020). However, they remain comparable to studies with similar maceration protocols (El Madany et al., 2023; Bouzenad, 2024). Our *C. fragile* extracts showed lower phenolic and flavonoid levels than reported elsewhere (Heffernan et al., 2015), while *Batophora* sp. had fewer phenolics than related green algae (Ruiz-Medina et al., 2022). For *P. tenerrima*, phenolic levels were in the same range as other *Laurencia complex* species (Bianco et al., 2015; Ruiz-Medina et al., 2022).

Antioxidant activities from our extracts ranged between 20–50% DPPH inhibition, lower than in some *Dictyota* and *Codium* studies (Kolsi et al., 2017; Imran et al., 2023) but higher than reported for *Palisada* species (Hmani et al., 2021). The general trend brown > green > red algae agree with previous observations (Kelman et al., 2012; Dang et al., 2018) and reflect differences in algae metabolite composition. Brown algae are rich in terpenes and pigments such as fucoxanthin, both with strong antioxidant properties (El Shoubaky and Salem, 2014; Sugawara et al., 2006). The relatively high antioxidant activity of *Batophora* sp. may relate to coumarins, previously described from dasycladales algae (Pérez-Rodríguez et al., 2001).

Regarding antifungal capacity, brown algae again showed the strongest inhibition of *Fusarium oxysporum* f. sp. *cubense* TR4, followed by moderate activity in the green algae *C. fragile* and red algae *P. tenerrima*. Similar patterns have been attributed to phenolics and terpenoids in brown algae (Belattmania et al., 2016; Khan et al., 2017). In *R. okamurae*, activity may also involve acidic vacuolar compounds (García-Gómez et al., 2018). The Ceramiales, including *P. tenerrima*, produce halogenated sesquiterpenes and other terpenoids with known antifungal properties (Vairappan et al., 2008; Stein et al., 2011; Alarif et al., 2011). In *C. fragile*, diosgenin and saponins may explain antifungal activity (Rizwan et al., 2024). These results highlight the predominance of polar bioactive molecules, since methanolic extracts consistently outperformed ethyl acetate.

In general, environmental and seasonal factors may further explain differences with previous studies, as metabolite profiles shift with location, season, and growth phase (Floreto et al., 1993; Mannivanan et al., 2008; Trigui et al., 2013).

Altogether, these results identify several macroalgae from the southeastern Spanish coast as promising sources of antioxidant and antifungal compounds. In particular, the methanolic extract of *R. okamurae* showed the strongest inhibition of *Fusarium oxysporum* f. sp. *cubense* TR4, together with relevant phenolic content and antioxidant capacity. LC–MS/MS analysis further revealed a chemically diverse metabolite profile, including phenolic derivatives, oxylipin-type compounds and terpenoid-related structures that may contribute to the bioactivity found. Antifungal results open new opportunities to explore the potential of algae extract as an potential candidate for managing Fusarium wiltin banana. The antioxidant capabilities of the algae tested extracts also open new opportunities to improve the health of the plants. These findings highlight the potential of Mediterranean macroalgae as natural resources for antifungal and antioxidant applications, while also supporting the valorization of biomass from invaders such as *R. okamurae*.

## 5. Conclusion

Our algal extracts exhibited antioxidant activity, especially those from brown algae, where polar metabolites such as phenolics and flavonoids could play a major role. *R. okamurae, D. fasciola*, and *P. tenerrima* also inhibited *F. oxysporum* TR4, reinforcing their potential as a source of natural antifungal agents.

Among invasive species, *R. okamurae* stood out with the highest yields, phenolic content, antioxidant capacity, and antifungal activity, positioning this species as a promising candidate for applications in agriculture and biomedicine. In addition, *Batophora* sp. consistently showed strong antioxidant performance, supporting its relevance as a source of bioactive compounds.

In conclusion, exploiting invasive biomass—particularly *R. okamurae*—could simultaneously provide bioactive resources and help mitigate ecological impact on Mediterranean coasts.

## Author Contributions

All authors contributed equally to the conception, design, execution, analysis, and writing of this study. All authors have read and agreed to the published version of the manuscript.

## Conflicts of interest

The authors declare no conflict of interest

## Funding

This research was funded by PID2020–119734RB-I00 Project from the Spanish Ministry of Science and Cropsafe Project 101209410 from the EU CBE JU.

## Acknowledgements

The authors would like to thank Luis Vicente López Llorca for his valuable support, insightful discussions, and guidance throughout this study.

## Notes

### Competing Interest Statement

The authors have declared no competing interest.

