## Supplementary figures and tables for "Screening antifungal and antioxidant activity of macroalgae from SE Spain highlights the invader *Rugulopteryx okamurae*"

Supplementary material

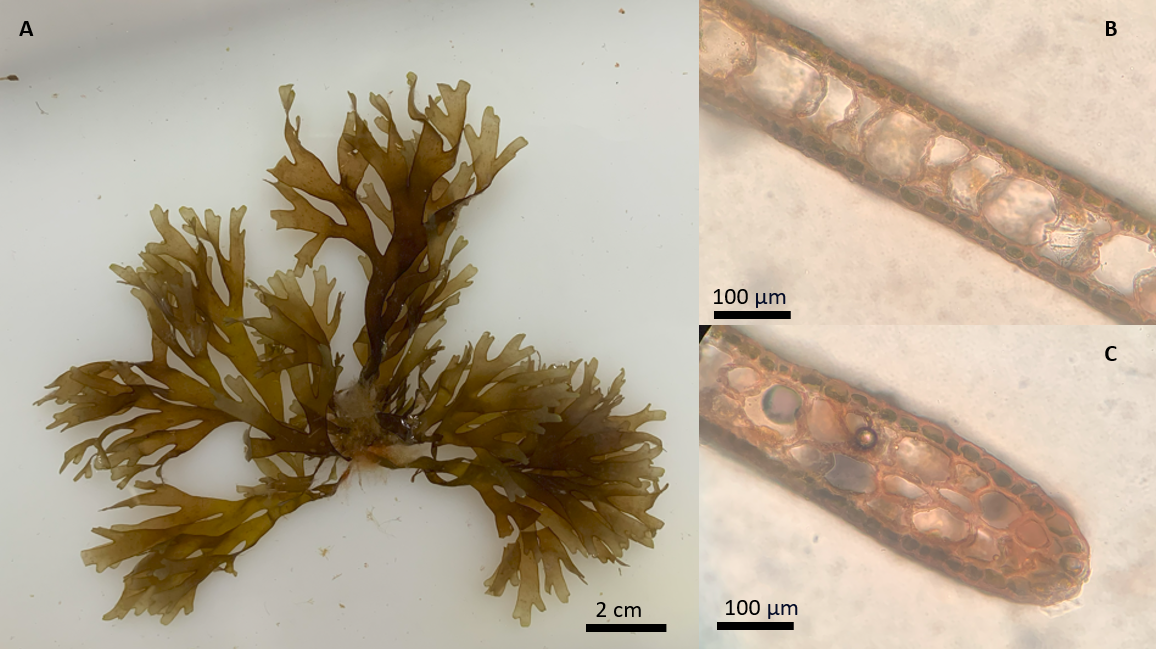

Figure S1. *Rugulopteryx okamurae*. General appearance of the alga (A); transverse section of the middle part showing the unistratified layer in the medial zone of the thallus (B); transverse section of the middle part showing the multistratified outer margin, with up to three cell layers (C).

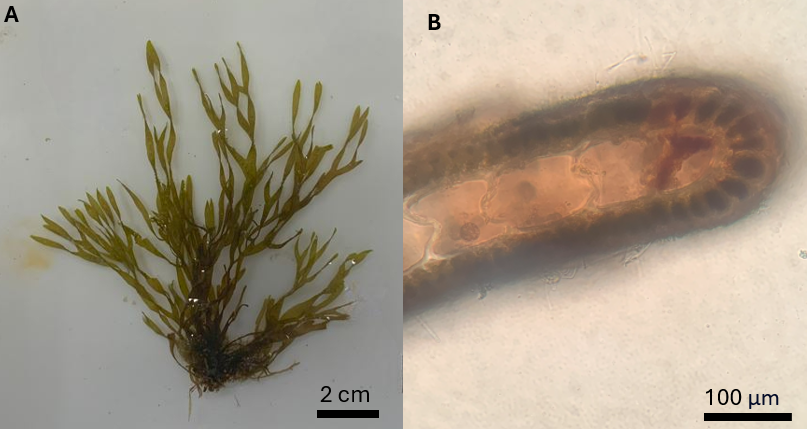

Figure S2. *Dictyota fasciola*. General appearance of the alga (A); transverse section of the middle part showing a unistratified medulla (B).

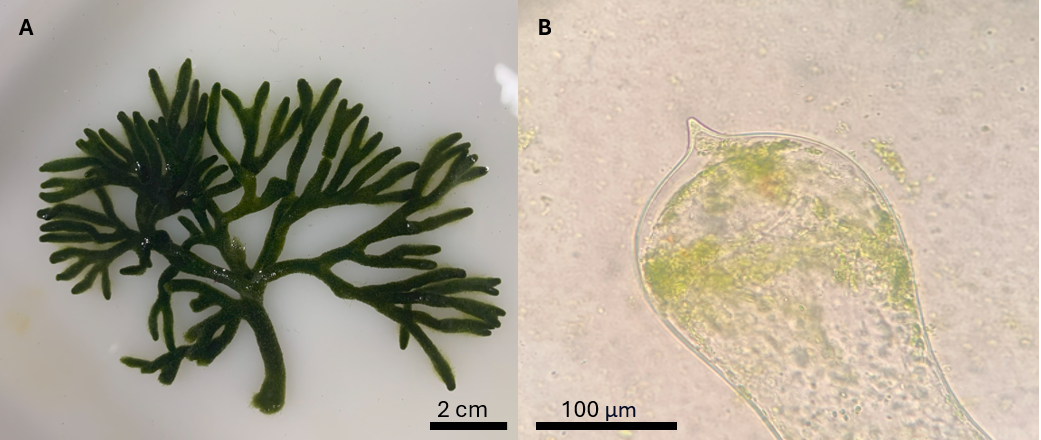

Figure S3. *Codium fragile*. General appearance of the alga (A); globose, mucronate utricles without corona (B).

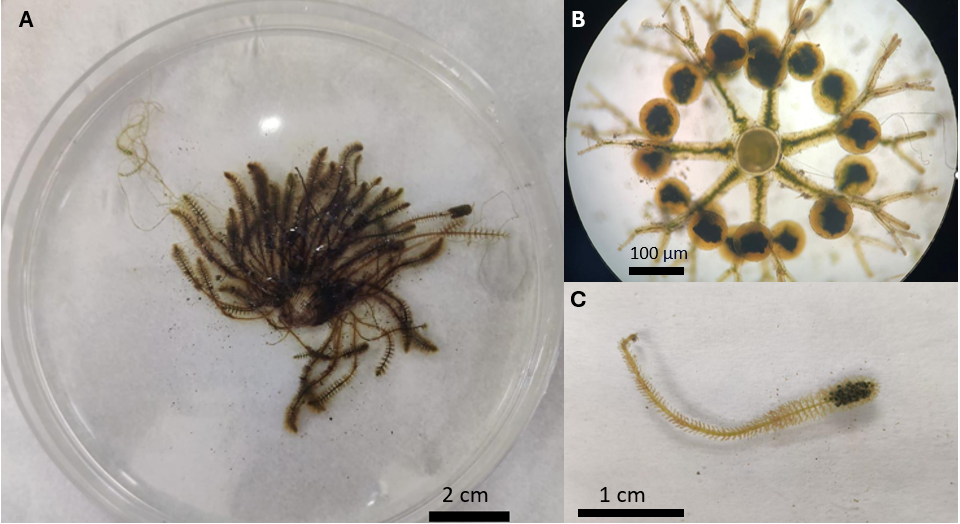

Figure S4. *Batophora* sp. General appearance of the alga (A); transverse section (B); detail of the branching pattern (C).

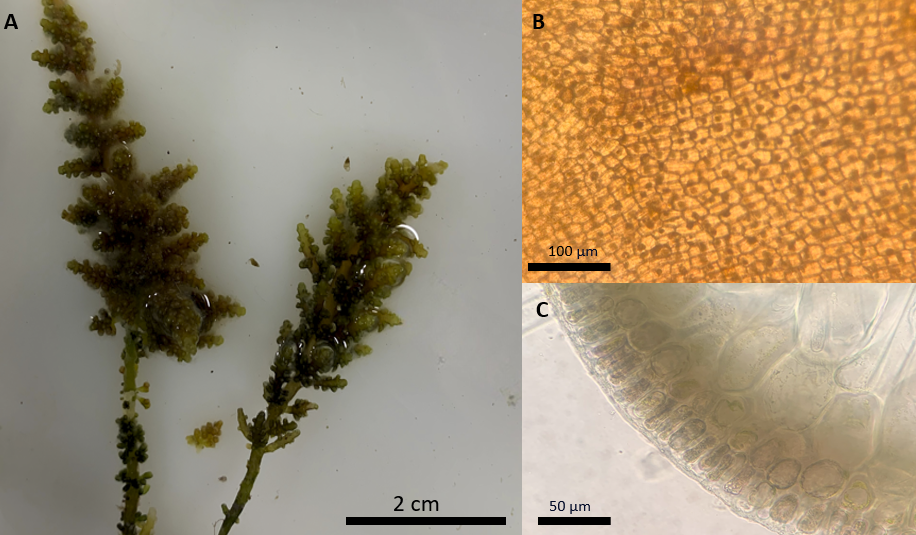

Figure S5. *Palisada tenerrima*. General appearance of the alga (A); surface view of the cells (B); transverse section of the middle part (C).

Table S1. Molecular identification of algal species included in this work. The top BLAST match is shown.

| Algae | Marker | Identity | *Accesion number* |
| --- | --- | --- | --- |
| *R. okamurae* | psbA | 98,73 % | MZ393490.1 |
|  | COI | 100 % | GQ425120.1 |
| *D. fasciola* | psbA | 97 % | MW225012.1 |
| *Batophora* sp. | rbcL | 99,22 % | MH54529.1 |
| *P. tenerrima* | COI | 90,57 % | MG030786.1 |

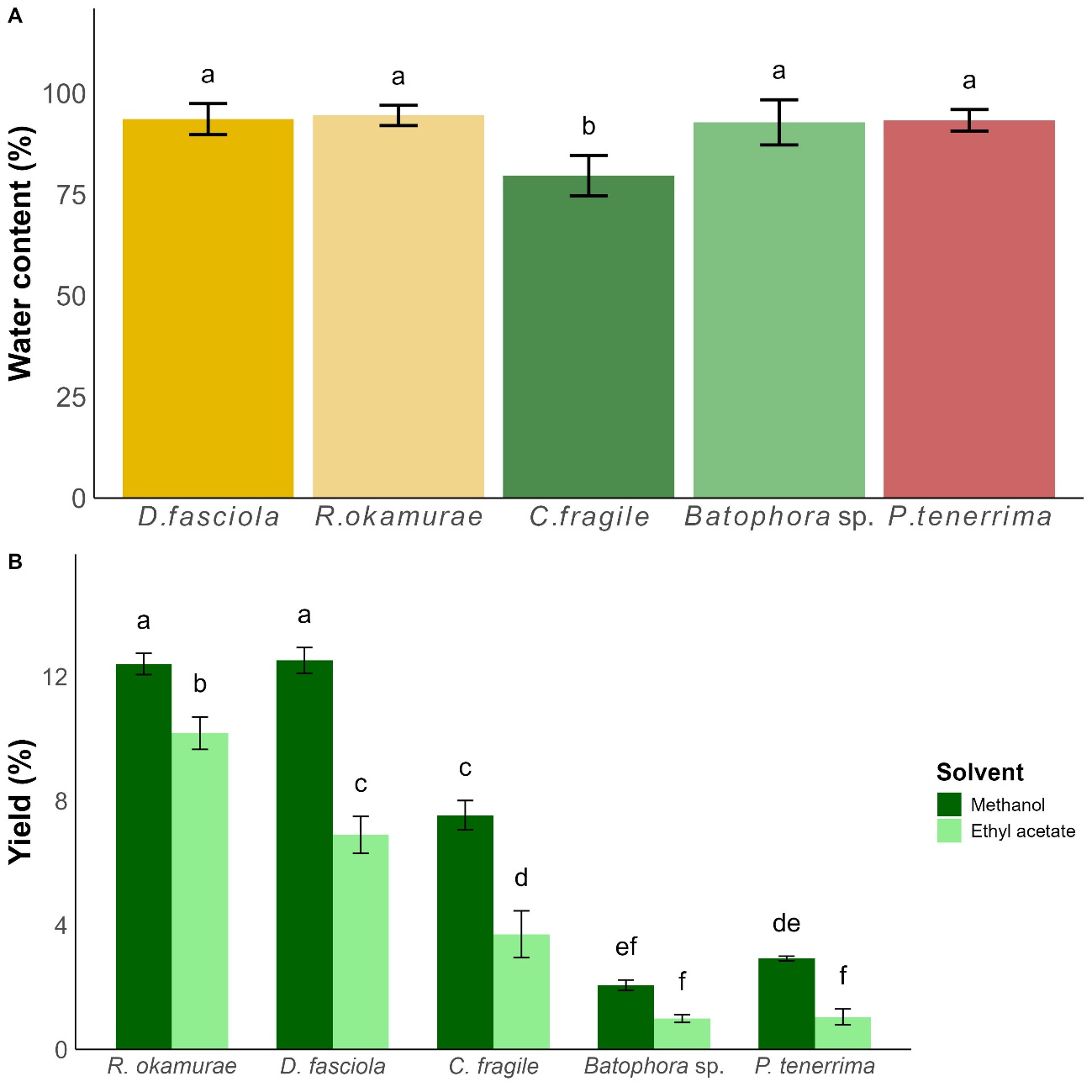

Figure S6. Characteristics of algal extracts used in this study. (A) Water content (%) of the different algae studied and (B) yield (%). Bars labeled with different letters indicate statistically significant differences (*p* < 0.05) according to the post hoc multiple comparison test. Bars sharing the same letter do not differ significantly from each other.

Table S2. ANOVA to assess water content differences among the studied algae. Df: degrees of freedom. Sum sq: sum of squares. Mean sq: mean squares.

|  | Df | Sum sq | Mean sq | F value | p.value |
| --- | --- | --- | --- | --- | --- |
| Algae | 4 | 559.0 | 139.75 | 12.97 | **<0.0001** |
| Residual | 28 | 301.7 | 10.78 |  |  |

Table S3. ANOVA to assess yield variation according to algae species and solvent used. Df: degrees of freedom. Sum sq: sum of squares. Mean sq: mean squares.

|  | Df | Sum sq | Mean sq | F value | p.value |
| --- | --- | --- | --- | --- | --- |
| Algae (A) | 4 | 468.10 | 117.03 | 644.27 | **<0.0001** |
| Solvent (S) | 1 | 64.20 | 64.17 | 353.28 | **<0.0001** |
| A x S | 4 | 19.60 | 4.90 | 29.97 | **<0.0001** |
| Residual | 20 | 3.60 | 0.18 |  |  |

Table S4. ANOVA to assess phenolic content variation according to algae species and solvent used. Df: degrees of freedom. Sum sq: sum of squares. Mean sq: mean squares.

|  | Df | Sum sq | Mean sq | F value | p.valor |
| --- | --- | --- | --- | --- | --- |
| Algae (A) | 4 | 8262 | 2066 | 31.39 | **<0.0001** |
| Solvent (S) | 1 | 3605 | 3605 | 54.78 | **<0.0001** |
| A x S | 4 | 4484 | 1121 | 17.04 | **<0.0001** |
| Residual | 50 | 3290 | 66 |  |  |

Table S5. ANOVA to assess flavonoid content variation according to algae species and solvent used. Df: degrees of freedom. Sum sq: sum of squares. Mean sq: mean squares.

| Término | Df | Sum sq | Mean sq | F value | p.valor |
| --- | --- | --- | --- | --- | --- |
| Algae (A) | 4 | 4482 | 1121 | 66.99 | **<0.0001** |
| Solvent (S) | 1 | 3548 | 3548 | 212.12 | **<0.0001** |
| A x S | 4 | 648 | 162 | 9.68 | **<0.0001** |
| Residual | 50 | 836 | 17 |  |  |

Tabla S6: ANOVA to evaluate antioxidant activity (% DPPH· radical inhibition) as a function of algal species, solvent used, and concentration applied. Df: degrees of freedom. Sum sq: sum of squares. Mean sq: mean square.

| Término | Df | Sum sq | Mean sq | F value | p.valor |
| --- | --- | --- | --- | --- | --- |
| Algae (A) | 5 | 60648 | 12130 | 834.01 | **<0.001** |
| Solvent (S) | 1 | 1824 | 1824 | 125.42 | **<0.001** |
| Concentration (C) | 3 | 858 | 286 | 19.67 | **<0.001** |
| A x S | 5 | 2919 | 584 | 40.14 | **<0.001** |
| A x C | 15 | 660 | 44 | 3.03 | **<0.001** |
| S x C | 3 | 97 | 32 | 2.23 | 0,08 |
| A x S x C | 15 | 671 | 45 | 3.07 | **<0.001** |
| Residual | 96 | 1396 | 15 |  |  |

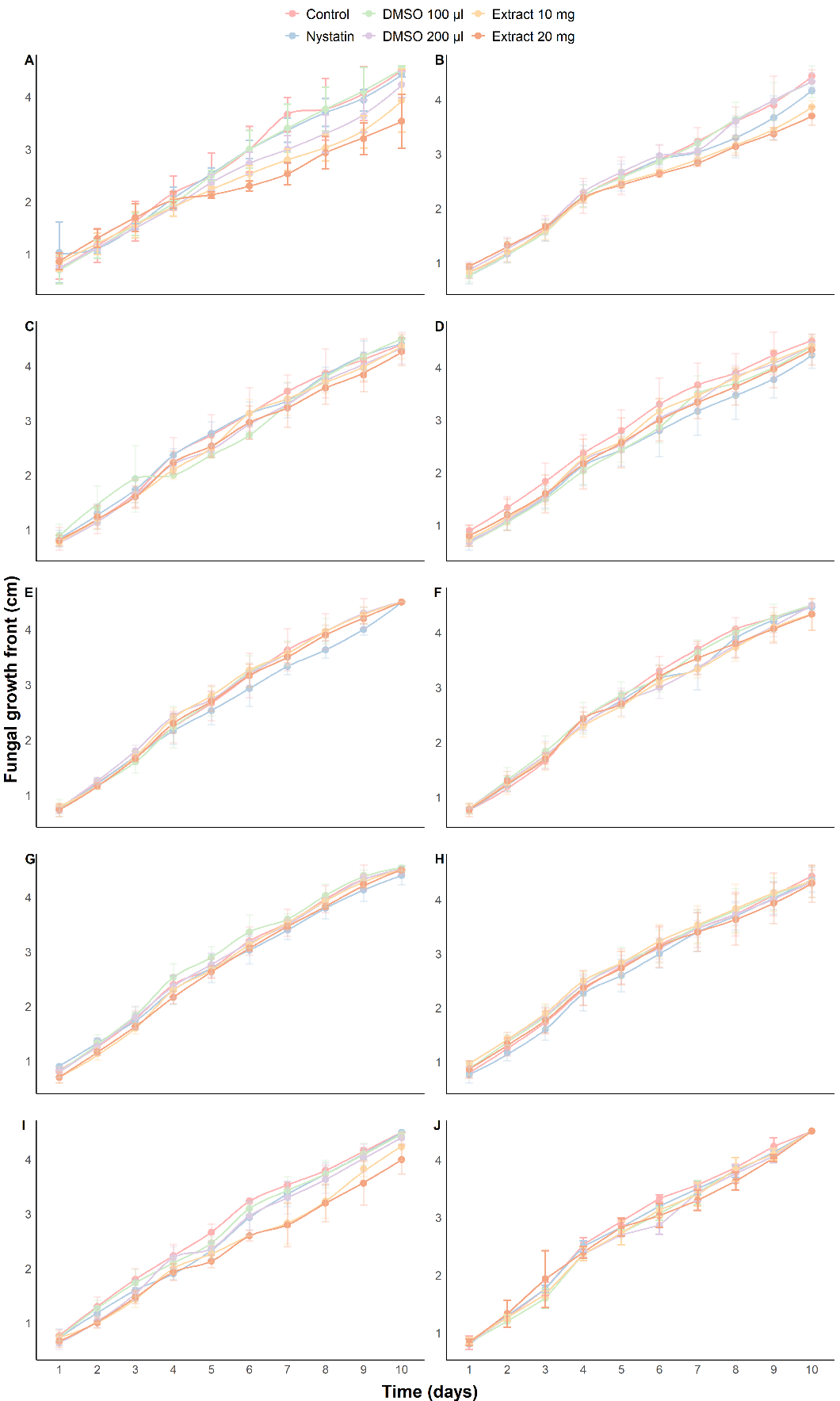

Figure S7: Effect of algal extracts obtained with methanol (left column) and ethyl acetate (right column) on the growth of banana wilt fungal pathogen *Fusarium oxysporum* f. sp. *cubense* Tropical Race 4 (FOC TR4). Panels correspond to extracts from *R. okamurae* (A–B), *D. spiralis* (C–D), *C. fragile.* (E–F), *Batophora sp.* (G–H), and *P. tenerrima* (I–J). Statistically significant differences (p < 0.05) are indicated by different letters.

Table S7: Type III analysis of variance using the Satterthwaite method for the linear mixed-effects model. The model evaluates the effect of treatment, day, and their interaction on fungal colony growth, with replication included as a random effect. **Sum sq**: sum of squares. **Mean sq**: mean squares. **NumDf**: numerator degrees of freedom. **DenDf**: denominator degrees of freedom.

|  | Sum sq | | | | Mean Sq | | | NumDf | | | | DenDf | | F.value | p.value |
| --- | --- | --- | --- | --- | --- | --- | --- | --- | --- | --- | --- | --- | --- | --- | --- |
| *R. okamurae* methanol extract | | | | | | | | | | | | | | | |
| Treatment(T) | 0.40 | | | | 0.08 | | | 5 | | | | 118 | | 10.68 | **<0.001** |
| Day (D) | 23.18 | | | | 2.57 | | | 9 | | | | 118 | | 338.98 | **<0.001** |
| T x D | 0.54 | | | | 0.01 | | | 45 | | | | 118 | | 1.58 | **0.02** |
| *R. okamurae* ethyl acetate extract | | | | | | | | | | | | | | | |
| T | | | 0.13 | 0.02 | | 5 | | | | 118 | | | 8.14 | | **<0.001** |
| D | | | 21.64 | 2.40 | | 9 | | | | 118 | | | 744.76 | | **<0.001** |
| T x D | | | 0.18 | 0.004 | | 45 | | | | 118 | | | 1.29 | | 0.13 |
| *D. fasciola* methanol extract | | | | | | | | | | | | | | | |
| T | 0.06 | | | | 0.01 | | 5 | | | | 118 | | 2.80 | | **0.01** |
| D | 25.65 | | | | 2.85 | | 9 | | | | 118 | | 627.08 | | **<0.001** |
| T x D | 0.14 | | | | 0.003 | | 45 | | | | 118 | | 0.71 | | 0.90 |
| *D. fasciola* ethyl acetate extract | | | | | | | | | | | | | | | |
| T | 0.05 | | | | 0.01 | | 5 | | | | 118 | | 5.32 | | **<0.001** |
| D | 28.96 | | | | 3.21 | | 9 | | | | 118 | | 1569.49 | | **<0.001** |
| T x D | 0.05 | | | | 0.001 | | 45 | | | | 118 | | 0.56 | | 0.98 |
| *C. fragile* methanol extract | | | | | | | | | | | | | | | |
| T | | 0.05 | | | 0.10 | | | | 5 | | | 118 | 4.75 | | **<0.001** |
| D | | 252.00 | | | 28.00 | | | | 9 | | | 118 | 1291.81 | | **<0.001** |
| T x D | | 0.74 | | | 0.01 | | | | 45 | | | 118 | 0.76 | | 0.85 |
| *C. fragile* ethyl acetate extract | | | | | | | | | | | | | | | |
| T | | 0.05 | | | 0.10 | | | | 5 | | | 118 | 4.75 | | **<0.001** |
| D | | 252.00 | | | 28.00 | | | | 9 | | | 118 | 1291.81 | | **<0.001** |
| T x D | | 0.74 | | | 0.01 | | | | 45 | | | 118 | 0.76 | | 0.85 |
| *Batophora* sp. methanol extract | | | | | | | | | | | | | | | |
| T | 0.07 | | | | 0.01 | | 5 | | | | 118 | | 8.40 | | **<0.001** |
| D | 28.08 | | | | 3.12 | | 9 | | | | 118 | | 1692.06 | | **<0.001** |
| T x D | 0.07 | | | | 0.001 | | 45 | | | | 118 | | 0.88 | | 0.66 |
| *Batophora* sp. ethyl acetate extract | | | | | | | | | | | | | | | |
| T | 0.07 | | | | 0.01 | | 5 | | | | 118 | | 16.86 | | **<0.001** |
| D | 23.83 | | | | 2.64 | | 9 | | | | 118 | | 2887.88 | | **<0.001** |
| T x D | 0.05 | | | | 0.001 | | 45 | | | | 118 | | 1.39 | | 0.07 |
| *P. tenerrima* methanol extract | | | | | | | | | | | | | | | |
| T | 0.44 | | | | 0.08 | | 5 | | | | 118 | | 24.68 | | **<0.001** |
| D | 27.21 | | | | 3.02 | | 9 | | | | 118 | | 836.82 | | **<0.001** |
| T x D | 0.13 | | | | 0.003 | | 45 | | | | 118 | | 0.84 | | 0.72 |
| *P. tenerrima* ethyl acetate extract | | | | | | | | | | | | | | | |
| T | 0.03 | | | | 0.006 | | 5 | | | | 118 | | 3.74 | | **<0.01** |
| D | 25.94 | | | | 2.88 | | 9 | | | | 118 | | 1673.75 | | **<0.001** |
| T x D | 0.07 | | | | 0.001 | | 45 | | | | 118 | | 0.92 | | 0.60 |

Figure S8: ANOVA assessing the effect of treatments of the different extracts on the growth front of FOC TR4. Df: degrees of freedom. Sum sq: sum of squares. Mean sq: mean squares.

|  | Df | Sum sq | | Mean sq | | F value | | p.value | |
| --- | --- | --- | --- | --- | --- | --- | --- | --- | --- |
| *R. okamurae* methanol extract | | | | | | | | | |
| Treatment (T) | 5 | 2.69 | | 0.53 | | 6.22 | | **0.004** | |
| Residual | 12 | 1.04 | | 0.08 | |  | |  | |
| *R. okamurae* ethyl acetate extract | | | | | | | | | |
| T | 5 | | 0.37 | | 0.07 | | 2.95 | | 0.057 |
| Residual | 12 | | 0.30 | | 0.02 | |  | |  |
| *D. fasciola* methanol extract | | | | | | | | | |
| T | 5 | | 0.16 | | 0.03 | | 0.47 | | 0.79 |
| Residual | 12 | | 0.84 | | 0.07 | |  | |  |
| *D. fasciola* ethyl acetate extract | | | | | | | | | |
| T | 5 | | 0.43 | | 0.13 | | 0.78 | | 0.57 |
| Residual | 12 | | 1.31 | | 0.10 | |  | |  |
| *C. fragile* methanol extract | | | | | | | | | |
| T | 5 | | 0.16 | | 0.03 | | 0.61 | | 0.69 |
| Residual | 12 | | 0.63 | | 0.05 | |  | |  |
| *C. fragile* ethyl acetate extract | | | | | | | | | |
| T | 5 | | 0.39 | | 0.07 | | 1.39 | | 0.29 |
| Residual | 12 | | 0.67 | | 0.05 | |  | |  |
| *Batophora* sp. methanol extract | | | | | | | | | |
| T | 5 | | 0.06 | | 0.01 | | 0.69 | | 0.63 |
| Residual | 12 | | 0.24 | | 0.02 | |  | |  |
| *Batophora* sp. ethyl acetate extract | | | | | | | | | |
| T | 5 | | 0.43 | | 0.08 | | 0.78 | | 0.57 |
| Residual | 12 | | 1.31 | | 0.10 | |  | |  |
| *P. tenerrima* methanol extract | | | | | | | | | |
| T | 5 | | 1.49 | | 0.29 | | 3.48 | | **0.03** |
| Residual | 12 | | 1.02 | | 0.08 | |  | |  |
| *P. tenerrima* ethyl acetate extract | | | | | | | | | |
| T | 5 | | 0.12 | | 0.02 | | 0.98 | | 0.46 |
| Residual | 12 | | 0.30 | | 0.02 | |  | |  |

Table S9: Multiobjective input matrix and Pareto classification for algal extracts (species × solvent). Values used in the 6-dimensional Pareto analysis (maximized criteria: antifungal inhibition at 10 and 20 mg, DPPH at 1 mg/ml, phenolics [µg GAE /mg], flavonoids [µg QE/mg], and yield [%]). “Pareto” marks non-dominated extracts (globally optimal across all criteria). Negative antifungal values indicate growth above the plate control on day 7 (no inhibition or slight stimulation). Abbreviations: MeOH, methanol; EtOAc, ethyl acetate.

| Specie | Solvent | Antifungal (10 mg) (%) | Antifungal (20 mg) (%) | DPPH (%) | Phenols  µg GA/mg | Flavonoids  µg QE/mg | Yield (%) | Pareto |
| --- | --- | --- | --- | --- | --- | --- | --- | --- |
| *R. okamurae* | MeOH | 23.30 | 30.50 | 44.7 | 62.3 | 35.0 | 12.4 | T |
| *D. fasciola* | MeOH | 3.62 | 8.39 | 33.8 | 45.5 | 44.6 | 12.5 | T |
| *Batophora* sp. | MeOH | -0.39 | 1.62 | 44.9 | 42.7 | 28.7 | 2.1 | T |
| *R. okamurae* | EtOAc | 10.10 | 12.10 | 25.5 | 16.3 | 18.4 | 10.2 | F |
| *D. fasciola* | EtOAc | 4.59 | 8.27 | 35.3 | 47.6 | 21.4 | 6.9 | F |
| *Batophora* sp. | MeOH | -1.95 | 1.95 | 43.2 | 20.5 | 7.29 | 1.0 | F |
| *C. fragile* | EtOAc | 1.40 | 7.14 | 35.8 | 29.8 | 10.6 | 7.54 | F |
| *C. fragile* | EtOAc | 3.11 | 8.11 | 30.2 | 21.0 | 4.24 | 3.71 | F |
| *P. tenerrima* | MeOH | 19.80 | 20.70 | 35.4 | 13.7 | 28.2 | 2.93 | F |
| *P. tenerrima* | EtOAc | 4.60 | 7.41 | 34.6 | 11.1 | 18.8 | 1.05 | F |

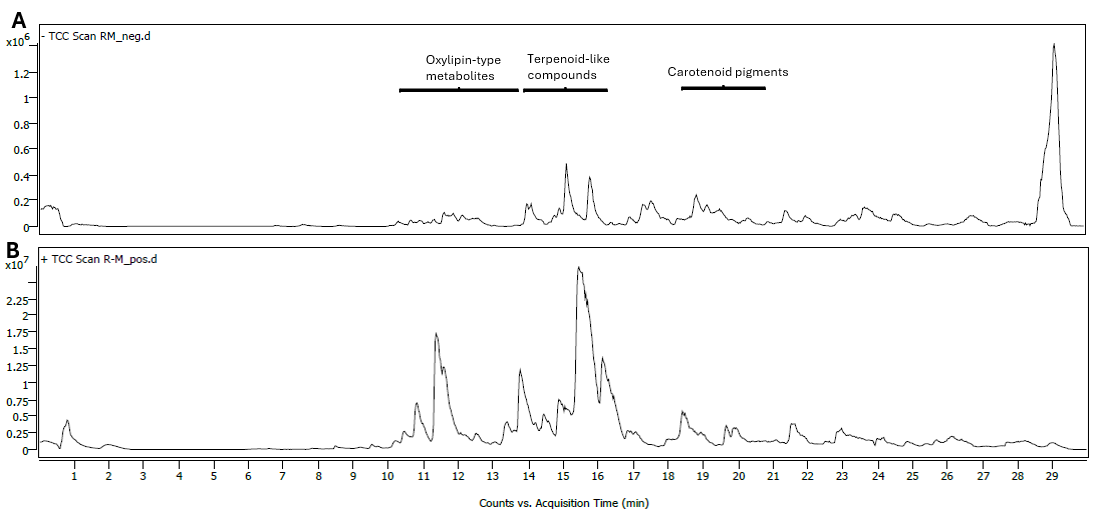

**Figure 10S.** LC–MS chromatographic profiles of the methanolic extract of *Rugulopteryx okamurae* obtained in negative (A) and positive (B) electrospray ionisation modes. Annotated regions indicate approximate retention time ranges where oxylipin-type metabolites, terpenoid-like compounds and carotenoid pigments were detected based on accurate mass and MS/MS fragmentation data.

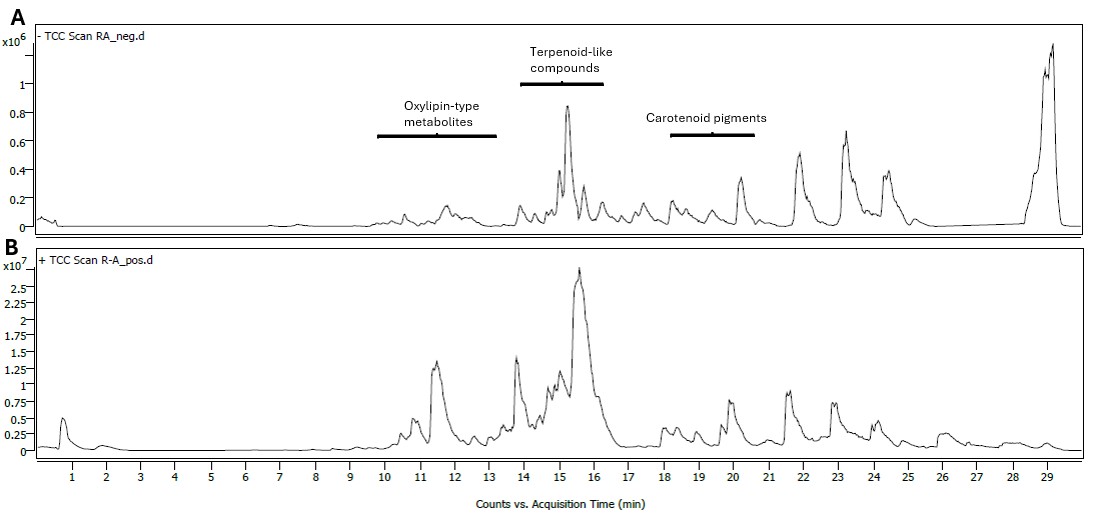

**Figure 11S.** LC–MS chromatographic profiles of the ethyl acetate extract of *Rugulopteryx okamurae* obtained in negative (A) and positive (B) electrospray ionisation modes. Annotated regions indicate approximate retention time ranges where oxylipin-type metabolites, terpenoid-like compounds and carotenoid pigments were detected based on accurate mass and MS/MS fragmentation data.
